# Sustainability of agricultural systems of indigenous people in Hidalgo, Mexico

**DOI:** 10.1101/2021.05.28.446198

**Authors:** Doris Leyva, Mayra de la Torre, Yaxk’in Coronado

**Affiliations:** Catedra-Conacyt-Unidad Regional Hidalgo, Centro de Investigacion en Alimentacion y Desarrollo A. C. Ciudad del Conocimiento y la Cultura de Hidalgo, Blvd. Santa Catarina S/N, San Agustin Tlaxiaca, Hidalgo, Mexico, 42163; Conacyt-Unidad Regional Hidalgo, Centro de Investigación en Alimentacion y Desarrollo A. C. Ciudad del Conocimiento y la Cultura de Hidalgo, Blvd. Santa Catarina S/N, San Agustin Tlaxiaca, Hidalgo, Mexico, 42163

**Keywords:** sustainability, MESMIS framework, rural agricultural systems, migrant remittances

## Abstract

Agricultural sustainability depends on complex relationships between environmental, economic and social aspects, in particular with the small farm holders from indigenous communities. This work was centered in two municipalities of Hidalgo State in Mexico, Ixmiquilpan (mainly irrigated systems) and El Cardonal (rainfed systems). Our objective was to understand the relationships between the small farm-holders and their agricultural systems. We evaluated the sustainability of their agricultural systems and did some recommendations. We applied the Framework for the Evaluation of Management Systems using Indicators (MESMIS, Spanish acronym), thirty-one indicators were identified, and the quantitative indexes were established to assess sustainability. The results showed that adaptability was a critical factor for irrigated and rainfed systems, the main problem identified was youth migration. Additionally, the access to water and economic resources, as well as environmental resources management, are imperious needs to increase the yield of agriculture crops. Therefore, a holistic approach taking into account the organization of small producers and synergy between indigenous knowledge and modern technologies, are required for the territorial development of the communities.

## 1. Introduction

The FAO defines sustainable agricultural development as “the management and conservation of the natural resource base, and the orientation of technological and institutional change in such a manner as to ensure the attainment and continued satisfaction of human needs for present and future generations. Such development… conserves land, water, plant and animal genetic resources, is environmentally non-degrading, technically appropriate, economically viable and socially acceptable.”[1]

The expansion of conventional agricultural techniques, mainly monoculture, and the massively increased use of agrochemicals cause an environmental crisis at global scale and raise the necessity of new approaches to solve it, such as sustainab1le agriculture. The main objectives of sustainable agriculture are: i) improving the health of producers and consumers (food security, organic agriculture); ii) maintain the stability of the environment (biological methods of fertilization and pest management); iii) ensure long-term benefits for farmers; and iv) considering the needs of current and future generations [2, 3]. Traditional farming systems could be an option, for example, rural indigenous communities in Central Mexico combine polyculture (corn, beans, squash, chili, and other crops) with organic fertilizers, no-tillage farming, no-agrochemicals and rainfall. Comparison of sustainability indicators of this traditional and other agricultural systems will allow to identify advantages and bottlenecks. Furthermore, socio-economic, and environmental indicators could serve as a tool in planning and decision-making processes at community and regional levels. These indicators belong to five main categories: production, resilience, adaptability, organization, social equity and self-sufficiency.

Concerns about agriculture sustainability go farther than environmental conditions and changes in internal trade; they also include the nature of the agricultural crisis that most of the countries are going through [4]. However, there are considerable discrepancies on translating the most appropriate philosophical and ideological aspects of sustainability, as well as what are the most appropriate methodologies to evaluate it. In this way, the evaluation of sustainability is affected by problems inherent to the multidimensionality of the concept itself, which includes ecological, economic, social, cultural, and temporal dimensions. Therefore, the evaluation requires a holistic [5] and systemic approach, where multi-criteria analysis predominates. Thus, there is a need to develop methods for evaluating the sustainability and performance of agricultural systems, as well as guide actions and policies for the sustainable management and preservation of natural resources. Even more Sustainable Management of Natural Resources (SMNR) has mainly focused on “sustainable agriculture”, but several authors have argued that SMNR should be understood in a broader sense, including activities such as forestry, livestock production, fisheries, mining, and ecotourism [6].

We consider the most adequate technology for evaluation of sustainability of complex agricultural systems is the framework for the evaluation of management systems using indicators (MESMIS, by its acronym in Spanish, Marco para la Evaluación de Sistemas de Manejo de Recursos Naturales Incorporando Indicadores de Sustentabilidad). This framework proposes six cyclic steps: 1) characterization of the systems; 2) identification of the critical points that are linked to the sustainability attributes, for example productivity, stability, reliability, resilience, adaptability, equity, and self-reliance; 3) identification and selection of indicators; 4) measurement and monitoring; 5) analysis and integration of data; and 6) conclusions and recommendations. [7–10]. Therefore, we used MESMIS, to evaluate and compare the sustainability of rainfed and irrigated agricultural systems of Otomies’ communities in two municipalities of the Hidalgo State in Mexico. These communities stand out as a low agricultural productivity region with semiarid climate and volcanic soil, hillocks, ravines, thorny scrub, and few streams. In the last five years the average annual precipitation was 140 milliliters in El Cardonal and 142 milliliters in Ixmiquilpan [11, 12]. In both municipalities, an ancestral polyculture system, referred to locally as ‘milpa’, is the basis of rainfed agriculture, and includes corn, beans, squash, chili, and many other crops. In irrigated lands, in contrast, monoculture systems are typical. This region was selected due to its long tradition of polyculture farming, and the presence of both irrigation and rainfed farming.

## 2. Materials and Methods

Mexico’s indegenous polulation is one of the two largets in the Americas, more than one in ten Mexicans speaks an indigenuos lenguage, and 5.7 percent of them live in Hidalgo State[13]. Our study area included 18 indigenous (Otomi or Hñähñu, as they refer to themselves) villages located in the municipalities of Ixmiquilpan and El Cardonal, in the Hidalgo State, Mexico (Figure 1). Ixmiquilpan is located at 20° 29’ 03” N, 99° 13’ 08” W, with an altitude of 1680 meters above sea level. This is an area dedicated to irrigated agriculture, the main crops being forage, vegetables, and grains [14]; the water source for irrigation is wastewater from Mexico City. El Cardonal is located between latitudes 20° 24’ 58” N and 20° 46’ 31” N and longitudes 98° 55’ 54” W and 99° 10’ 46” W, with an altitude between 900 and 2900 meters above sea level. This is an area dedicated to rainfed agriculture and grazing livestock (mainly sheep); the main crops are agave, corn, and fruit trees [15–17]. Inhabitants of both municipalities belong to the Otomi culture; their villages are communities of high and very high marginalization, and medium social backwardness.

**Figure 1.**
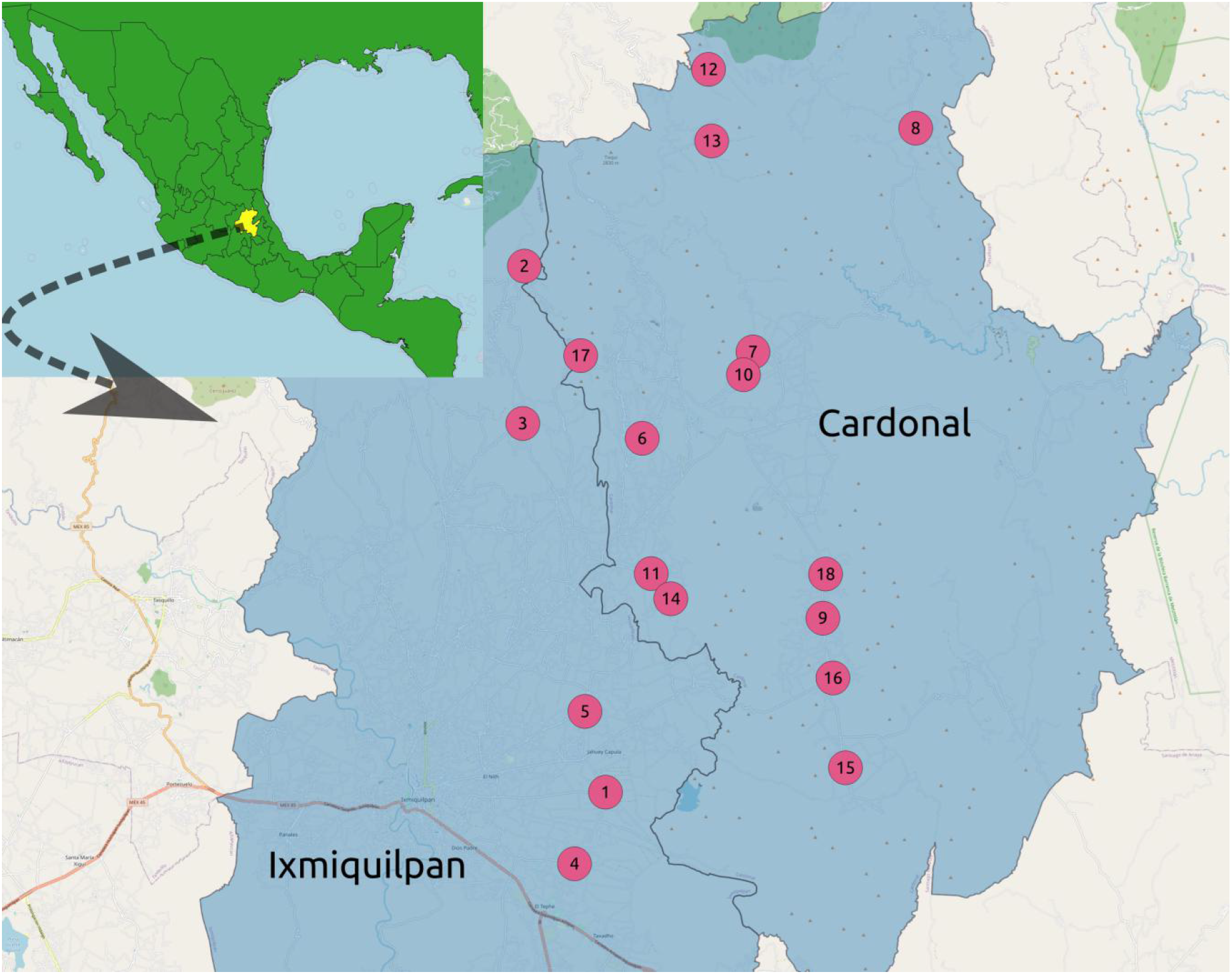
Location of municipalities and communities, **Ixmiquilpan**: (1) Bangandhó, (2) El Nogal, (3) El Olivo, (4) Pueblo Nuevo, (5) San Pedro Capula. **El Cardonal**: (6) Cerro Colorado, (7) Chalmita, (8) Cieneguilla, (9) Durango Daboxtha, (10) El Bondho, (11) El Botho, (12) El Potrero, (13) El Tixqui, (14) Los Reyes, (15) Pozuelos, (16) San Andrés Daboxtha, (17) San Miguel Jigui, (18) Santa Teresa Daboxtha.

We interviewed small farm holders who participated in the governmental program “Territorial Development Program” (PRODETER, by its acronym in Spanish) supported by the Ministry of Agriculture and Rural Development (SADER, by its acronym in Spanish) and live in the municipalities of Ixmiquilpan and El Cardonal. We adapted a mobile application to study the social, environmental, and economic relationships of agricultural systems. The mobile application koBoCollect v1.25.1, [18] allows collecting quantitative and qualitative data through a series of questions, including multimedia files, such as photos and GPS location. Data collection was carried out during January and February 2020, under a non-probability test sample [19, 20], considering between 15 and 20 percent of the producers registered in the database of PRODETER (121 interviews). Data includes socio-economic characteristics, family structure, economic incomes and outcomes from agricultural and non-agricultural activities, land use, food crops and yields, use of crops (sale, self-supply), agricultural practices, peasant awareness of climate change, use of chemical and biological pesticides, fertilizers, and herbicides, agricultural infrastructure, and adoption of new technologies. The product systems included in PRODETER are sheep, apple, and olive for Ixmiquilpan and corn, agave, sheep, and olive for El Cardonal. We selected those farmers that have corn crops and included in our study all the other agricultural products including crops, livestock, fruit trees, etc.

Following the MESMIS methodology, the first step was the characterization of the system including socio-economic and environmental features. The second step focused on identification of the critical points related to the attributes of productivity, stability, reliability, resilience, adaptability, equity, and self-reliance. The third stage was the selection of sustainable indicators and their corresponding normalization, which were flexible, easy to measure and to understand, and comprise social, economic, and environmental development (Table 1), selection of indicators followed the structure proposed by the MESMIS methodology; and each indicator was calculated according to its specific class and normalized to one hundred. The fourth was measurement and monitoring, followed by analysis and integration of data. The last stage was conclusions and recommendations.

**Table 1.**
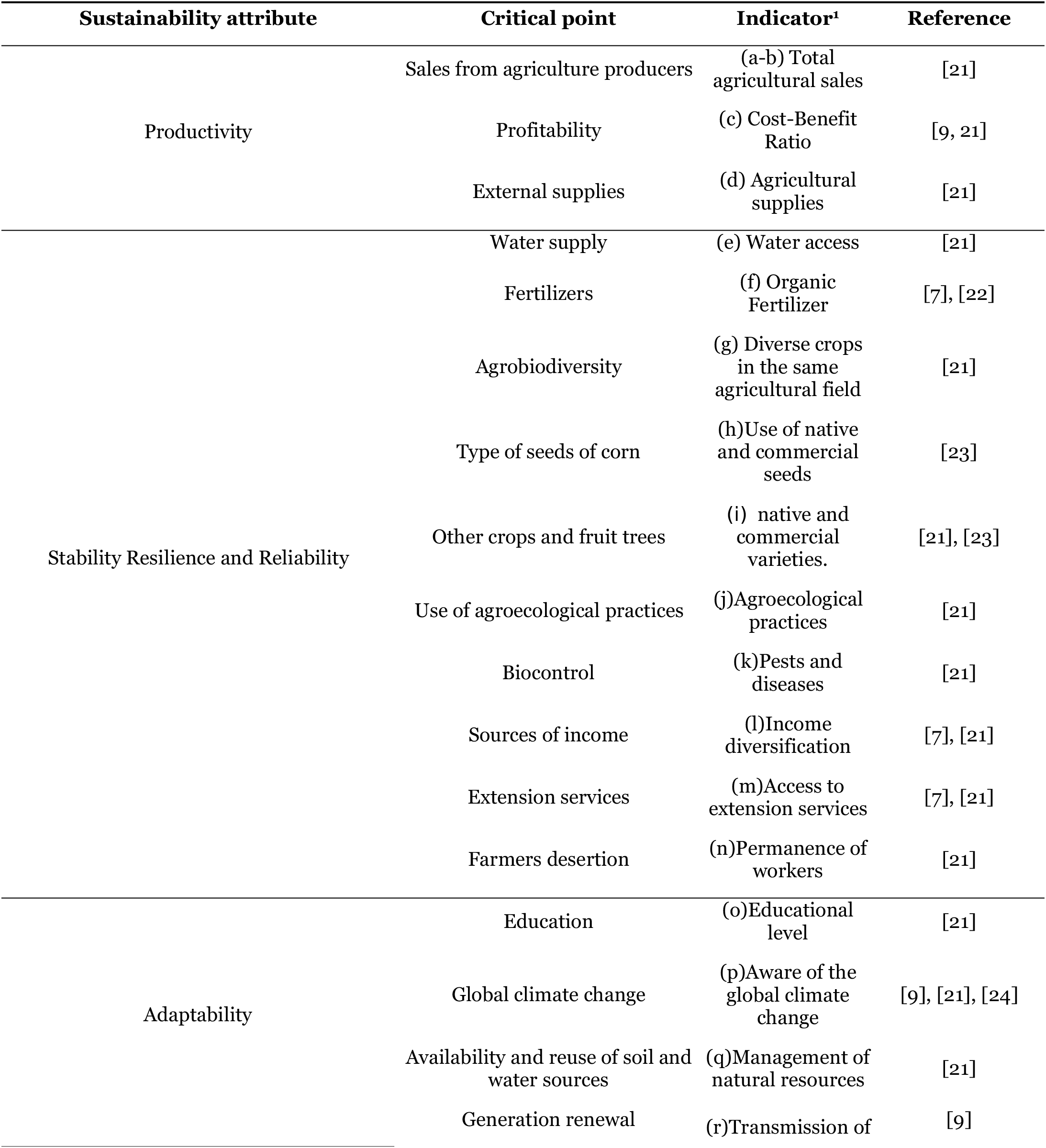
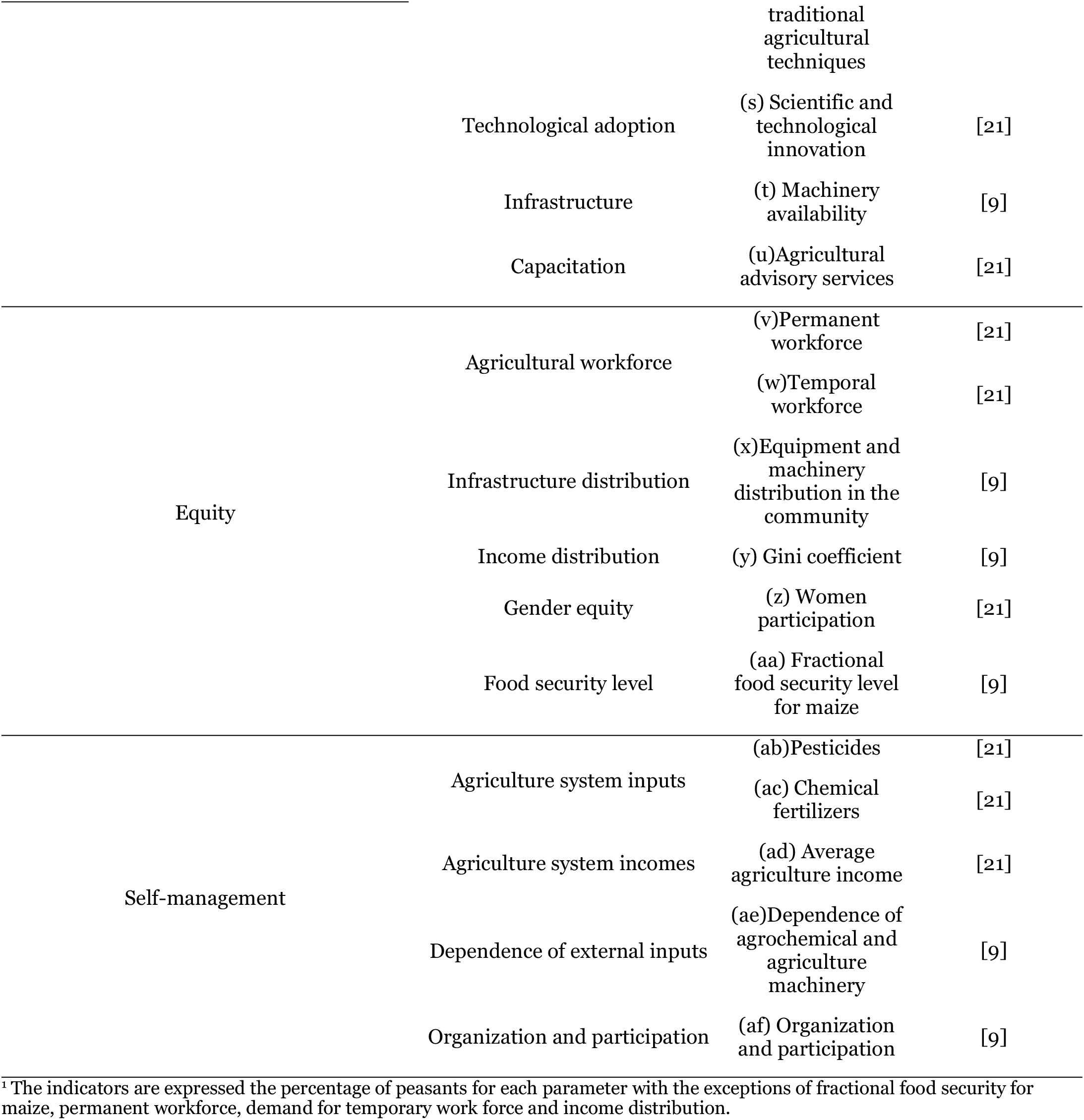
Sustainability attributes, critical points and indicators selected for the agricultural production systems.

For each indicator we define a parameter, classification, or calculation to quantify its representative value for sustainability. Each indicator was defined as follows:

a. Total agricultural sales for irrigation systems were calculated considering all the agricultural sales. The parameters used for classification were peasants with lower sales than the second quartile of the total peasants’ sales distribution, with sales between second and third quartile, and sales higher than the third quartile. The average yield was calculated based on the last year, either from irrigated or rainfed fields.
b. Total agricultural sales for rainfed systems were calculated as before described in (a)
c. Cost-Benefit Ratio (CBR) was calculated considering all the agricultural inputs divided by the total sales associated with agricultural activities for each producer.
d. Agricultural supplies were the average of consumables and capital inputs. Three levels were defined: lower than the first quartile, between the second and third quartile and greater than the third quartile.
e. Water access includes the natural water sources and irrigation channels available for agricultural activities. The parameters were defined as minimum when only rainfall was available; average when two different sources of water were accessible; and high when three or more sources of water were handy.
f. Three parameters were defined for fertilizers use only of organic fertilizer, employment of both organic and chemical fertilizers and no-fertilizers.
g. Agrodiversity was classified as low when it was monoculture farming, medium when two crops were intercropped, and high when more than two crops were intercropped.
h. For corn seeds the parameters were defined as only native seeds, commercial seeds, and a mix of native and commercial seeds.
i. For crops and fruit trees native or commercial varieties were considered and the definition is the same of (h). Backyard cultures where also included.
j. For agroecological practices the parameters were low when no agroecological practices were used, medium when only one practice was implemented and high when two or more practices were adopted.
k. For pests and diseases, the parameter was defined as high when agriculture pests and diseases were present in corn and intercropped crop, medium when only corn was affected, and low when they were not detected.
l. Income diversification was calculated as the number of incomes generating activities. It was defined as low when the income was only from agricultural activities, medium when the farmer realized other remunerated activity besides agriculture, and high when he or she had at least two more productive activities in addition to agriculture.
m. For access to extension services, the parameter was defined as high when peasants received technical assistance, low when they did not receive it but had requested it, and null when they did not receive assistance and they did not know with certainty if they need it.
n. For generational renewal, we considered workers as youths from 18 to 34 years, adults from 35 to 60 years and seniors older than 60 years.
o. Related to the educational level, inhabitants were classified in three levels according to their scholar level: uneducated, the ones that did not attend school; in the second level we included those who had elementary or junior high school studies, and in the third level the ones with high school or college studies.
p. For the impact of global climate change, we considered three groups of producers: those who were aware of global climate change and thought it had a big impact on their crops; a second group that thought global climate change did not affect their crops and a third group that did not know whether it had an effect.
q. Management of natural resources. Percentage of producers who cares about water and soil conservation. The parameter was classified as optimal if they practice water and soil conservation; medium, if they practice water or soil conservation, and non-conservation.
r. Generational transmission of knowledge was the percentage of producers who transmit knowledge and skills to younger. The parameters were, high, low, and null.
s. Technological adoption was considered as the percentage of producers who had access to cell phones, internet, etc. The parameters were, high, low and null.
t. Infrastructure was the percentage of farmer that have access to tractors and other farmer equipment.
u. Agricultural advisory service was defined as the percentage of farmers requiring information dissemination, training and advisory services.
v. Permanent workforce was the number of family members who were permanent agriculture-workers per hectare per year. The optimal indicator was fitted to 3 workers per hectare.
w. Demand for temporary workforce, was the number of hired agriculture workers per hectare per year. The optimal indicator was fitted to 3 workers per hectare.
x. Distribution of machinery and equipment was the percentage of producers who own tractors and farm equipment.
y. The Gini coefficient was used to assess inequality. It is defined as previously described [25, 26]:

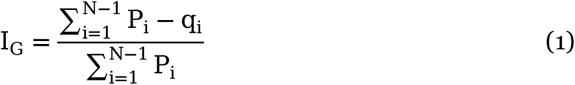

Where q_i_ is the accumulated relative income of the sample for each system and P_i_ is the cumulative relative frequency of the population. In our case qi was the peasant’s accumulated relative income by agricultural sales and Pi was the accumulated population, i.e., the number of members of the peasant’s family. The Gini coefficient was calculated for small farm holders owning either rainfed or irrigated farmlands.

z) The Participation of women was the percentage of farmer women.
aa) Chemical pesticides corresponded to the percentage of farmers who used chemical pesticides.
ab) Chemical fertilizers corresponded to the percentage of peasants who used chemical fertilizers.
ac) The Fractional Food Security Level (FFSL) for corn was calculated as [9]:

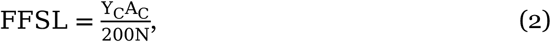

Where Yc is the corn yield in Kg/ha, Ac is the net corn-area cultivated and N is the number of farmers’ family members. Corn food security level is 200 Kg per Mexican per year [27].

ad) Total income was the average household income per farmer per year.
ae) Dependence of external inputs was the percentage of farmers who purchased fertilizers, herbicides, fungicides, and use agricultural machinery.
af) Organization was the percentage of farmers who are members of a farmers’ association.

## 3. Results

### 3.1. Characterization of the system

El Cardonal and Ixmiquilpan municipalities are mostly “Hñahñus” (Otomi) and speak Spanish and Hñahñu, they live in nuclear families, and only very few have extended families. The average age of farmer is 51 years and most of them have elementary studies; in contrast their offspring and relatives have an average age of 30 years and have junior and senior high-school studies [28]. Typically, the peasants’ farmland extensions are less than one hectare in Ixmiquilpan, and from 2 to 4 hectares in El Cardonal. Some farmers own and cultivate up to three different agricultural lands. In El Cardonal agricultural lands are rainfed while in Ixmiquilpan 95% of the lands have irrigation systems. These irrigation systems use residual water from Mexico City. It is worth mentioning that in Ixmiquipan most of the agricultural lands are communal; in contrast in El Cardonal 70% are private properties. Even when parcels are very small these are not rented to other producers. Most noteworthy, migration plays an essential role in both municipalities; inhabitants migrate to other Mexican States or mainly to the USA, to improve their living standard. In El Cardonal 62% of the families have at least one member who migrated and in Ixmiquilpan 44% of the families.

**Table 2.**
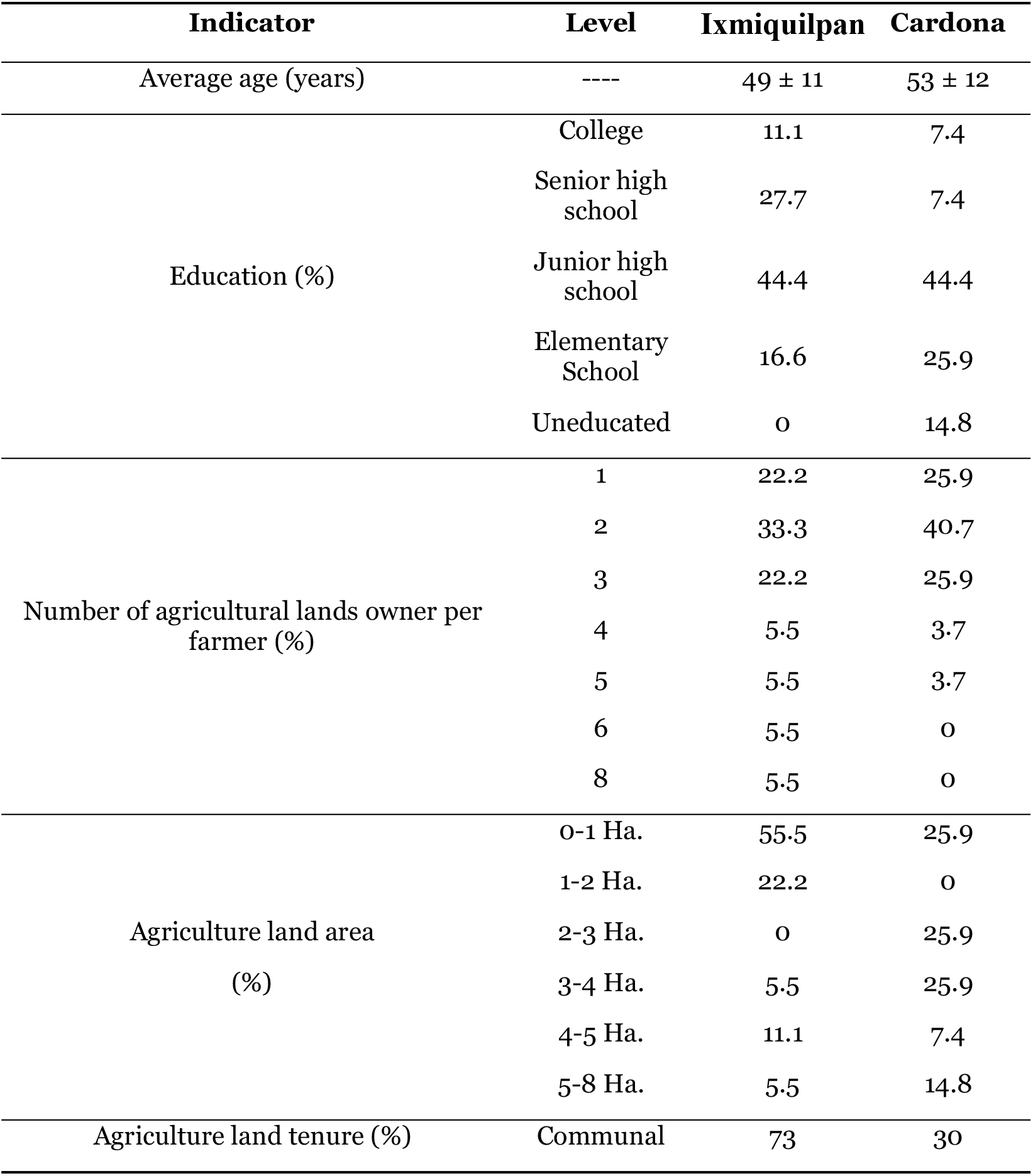
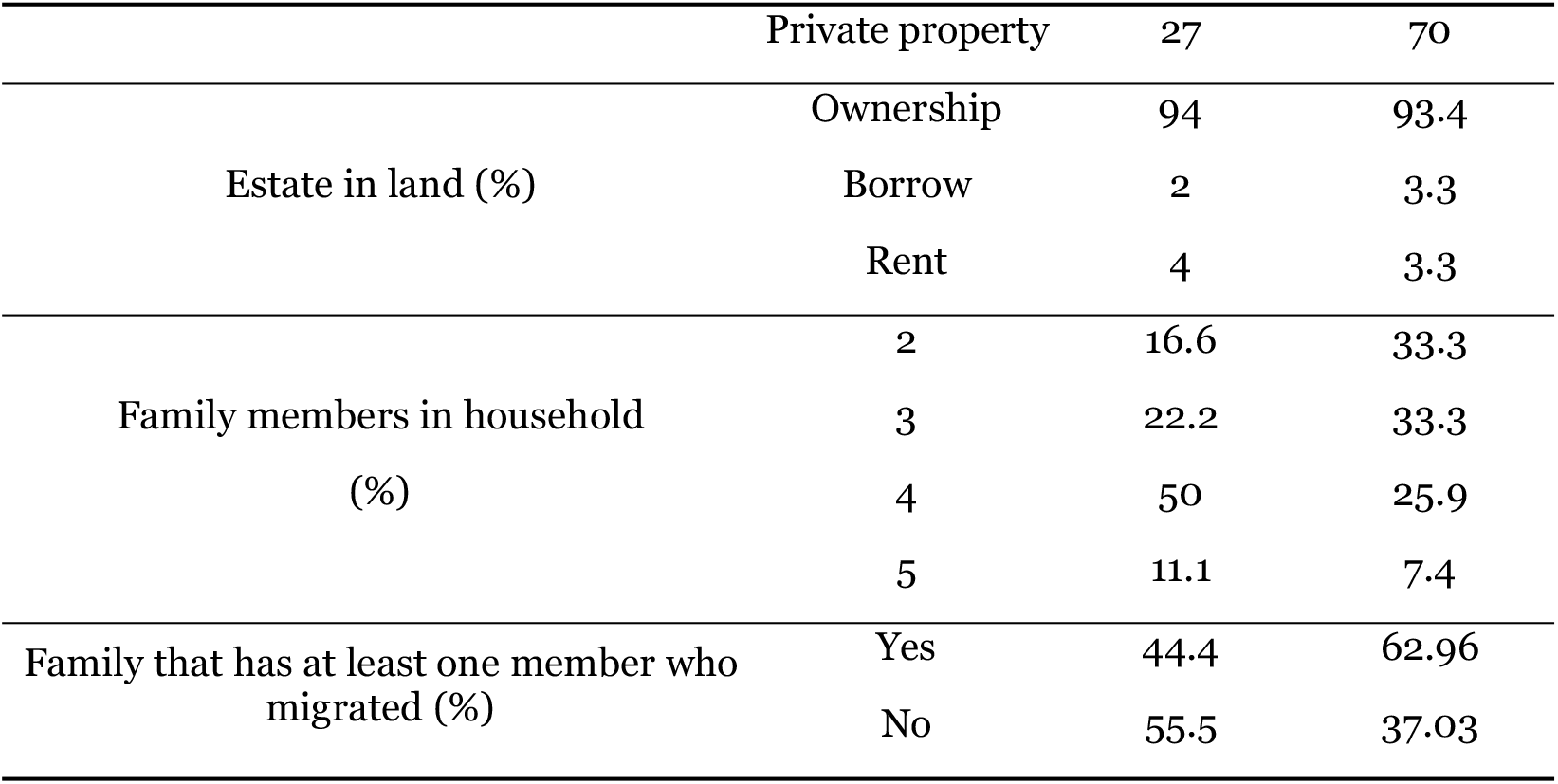
Socioeconomic indicators of El Cardonal and Ixmiquilpan municipalities.

### 3.2. Sustainability attributes and critical points

#### 3.2.1. Productivity

Productivity. The total agriculture-sales in the high class is higher for rainfed lands than the irrigated ones, although corn-yield is lower. Indeed, rainfed agricultural products include several crops, fruits, agave, livestock and edible wild plants and edible insects and backyard products. Notably, the inputs including investment and agricultural supplies for irrigated and rainfed system are similar (Table 3).

**Table 3.**
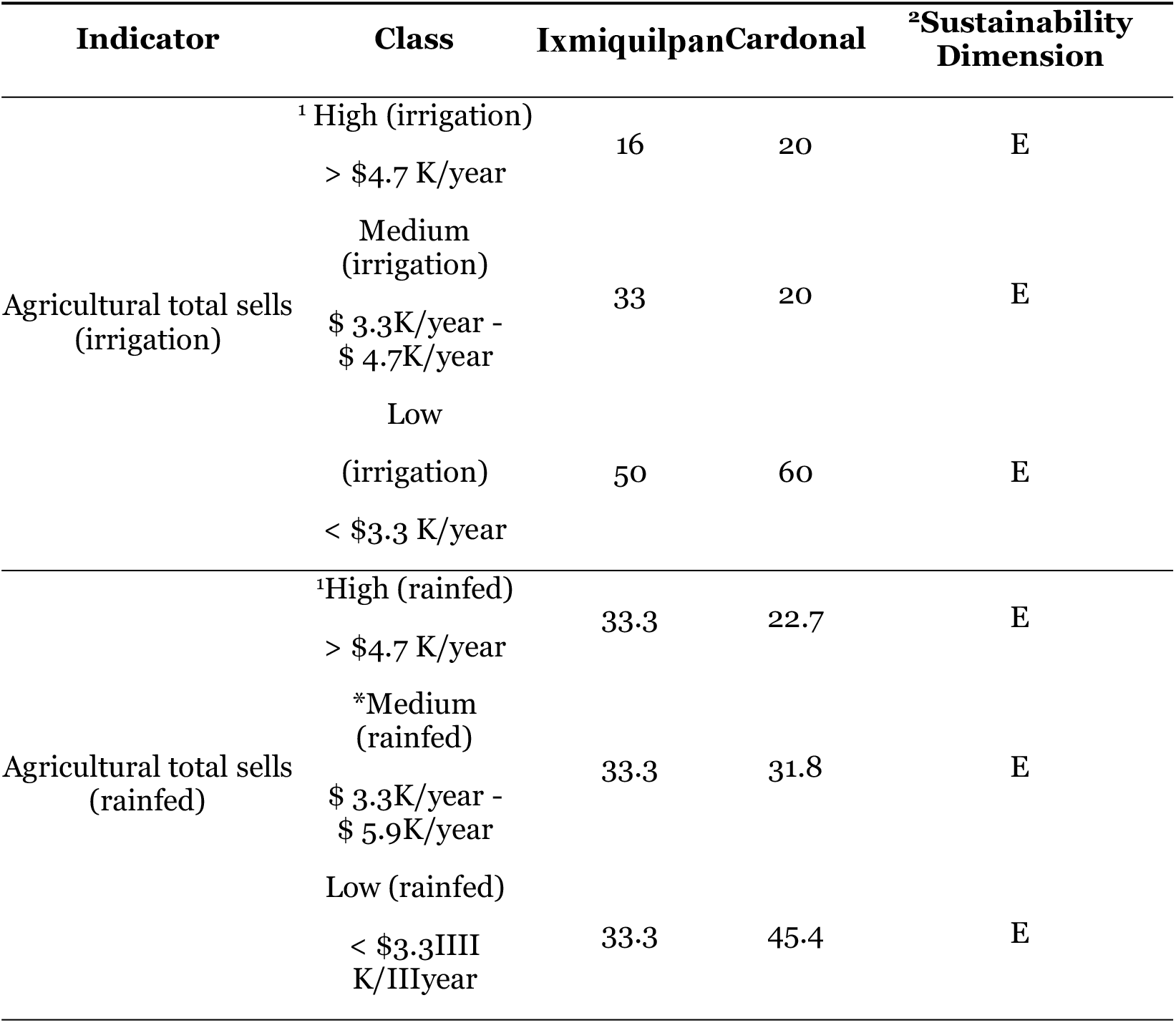
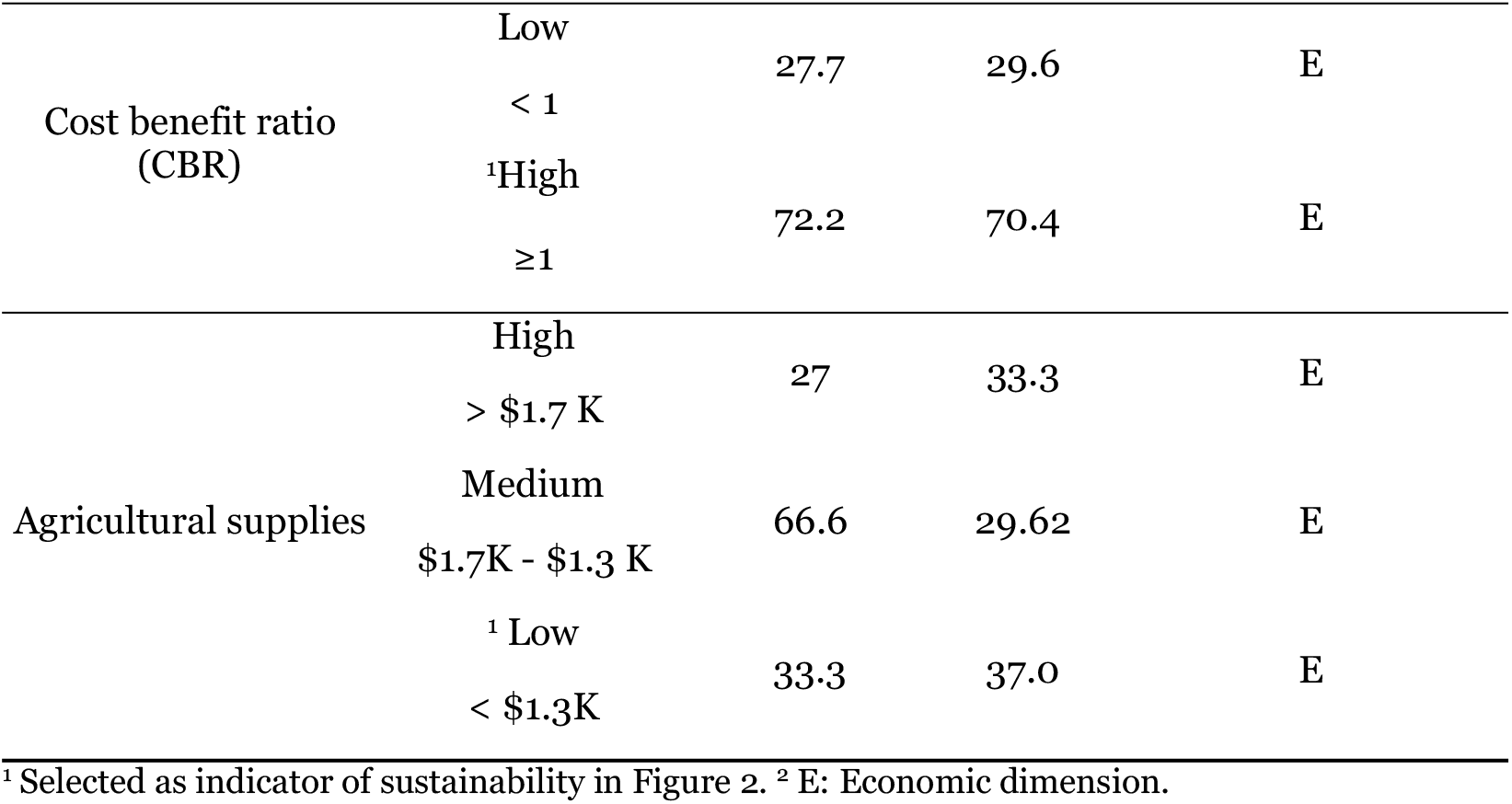
Productivity indicators for El Cardonal and Ixmiquilpan municipalities.

#### 3.2.2. Stability, resilience, and reliability

Water shortage, dry land, and drought, obligate farmers to develop strategies and actions for keeping their crops, such as polyculture-systems and use of natural fertilizers. Usually, native corn is grown in rainfed systems and backyard. Besides, in the irrigated farmlands hybrid and commercial seeds are used. It is worth mentioning that farmers in rainfed systems use agroecological practices, including minimum tillage, natural fertilizers, corn and intercropped crops and pest-repelling plants (Table 4). Notwithstanding pests and phytopathogens are an important problem mainly in the irrigated farmlands, however identification and biological control of pests and diseases in both agricultural systems could increase resilience.

**Table 4.**
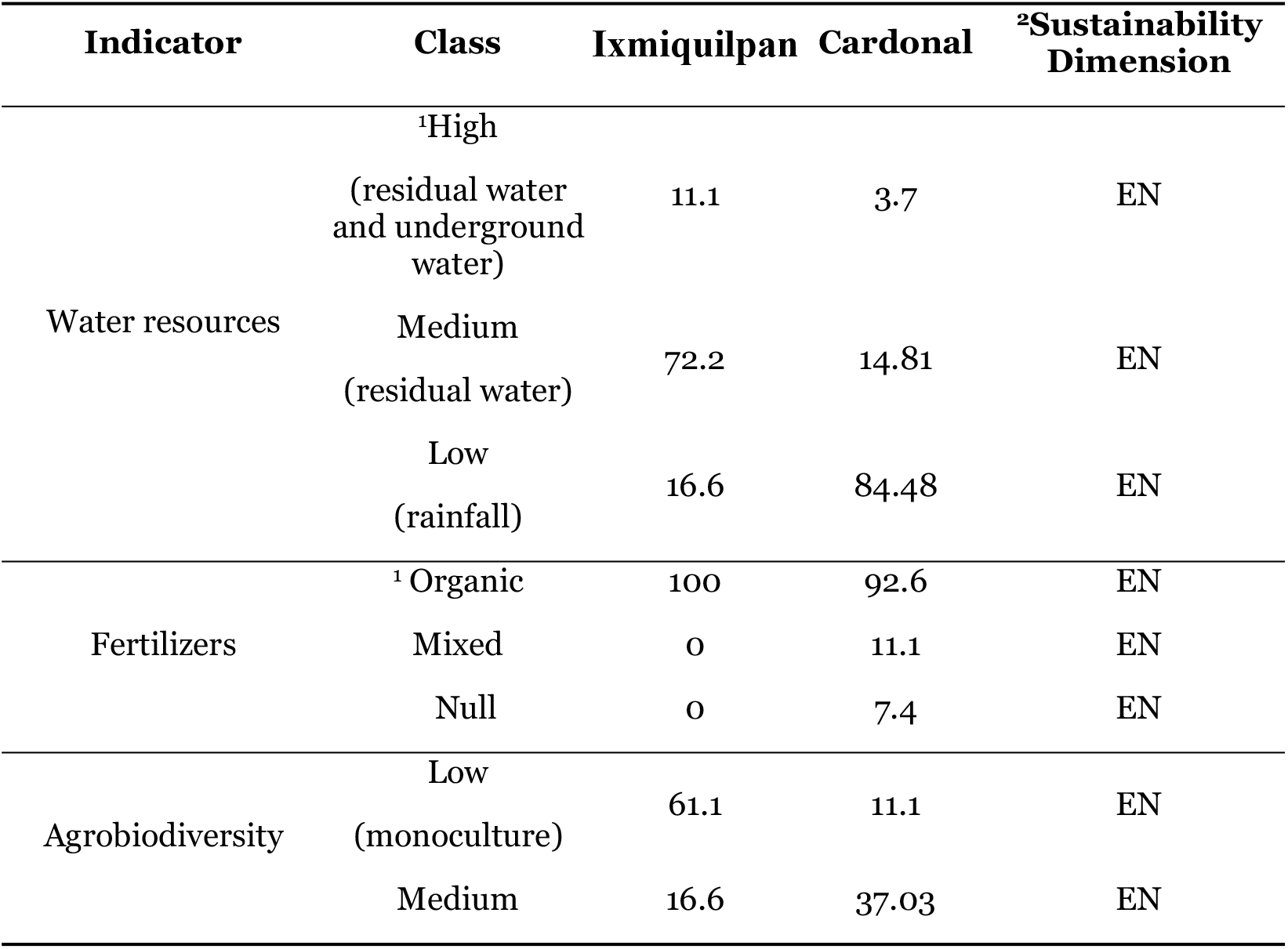
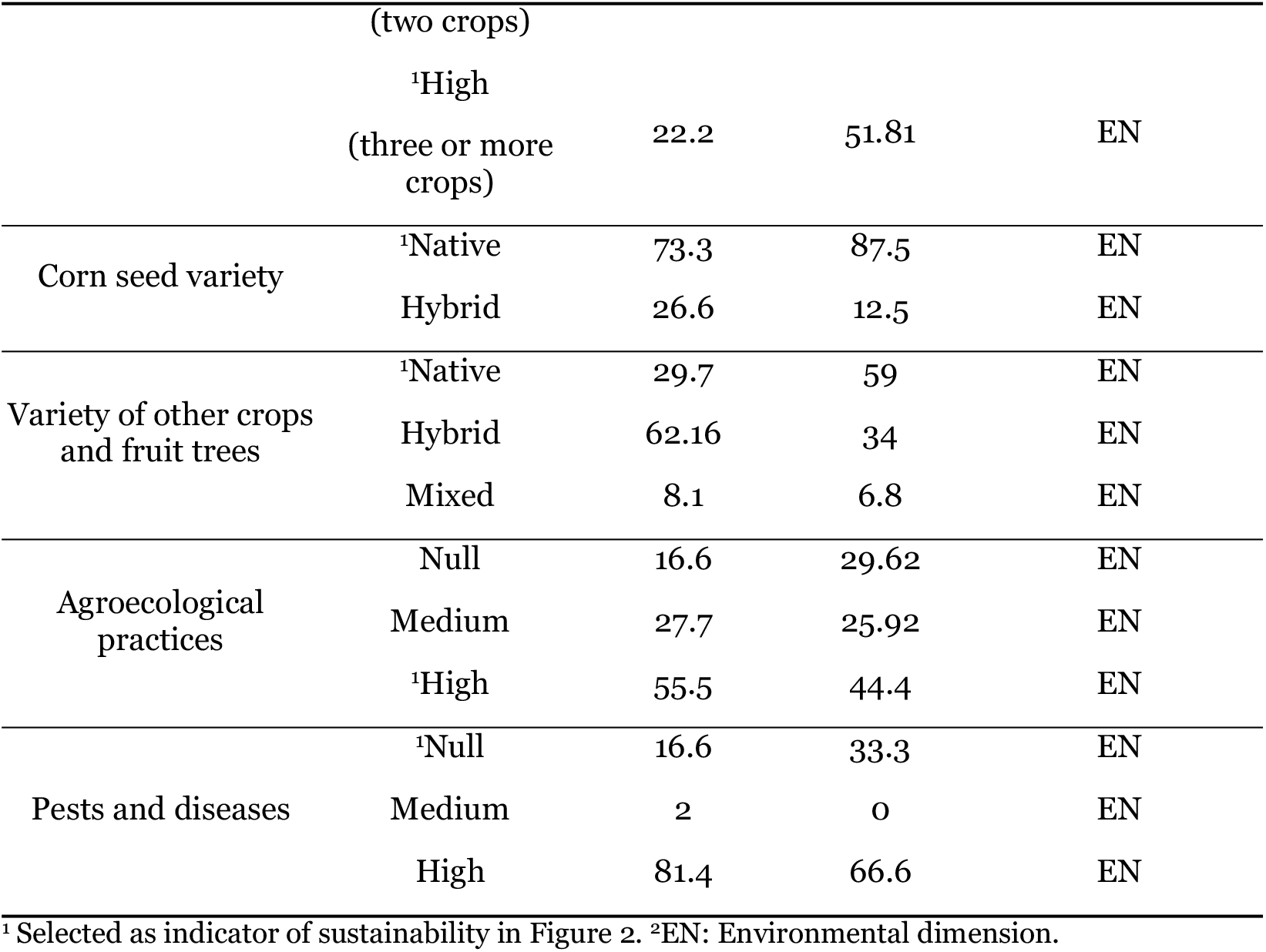
Stability, resilience, and reliability indicators for El Cardonal and Ixmiquilpan municipalities.

#### 3.2.3 Adaptability

This attribute is the main bottleneck for both municipalities. Although smallholder farmers have noticed a climate change, they are not aware and have not yet adopted some new strategies. Conservation of soil is a concern for few, but for all of them water supply is the most important problem. However, they do not have a plan for water management, neither for diversification of water sources. Even more, in both municipalities, traditional knowledge is scarcely transmitted to the new generations, and it is being lost, since the youths abandon both farmlands and their villages (Table 5).

**Table 5.**
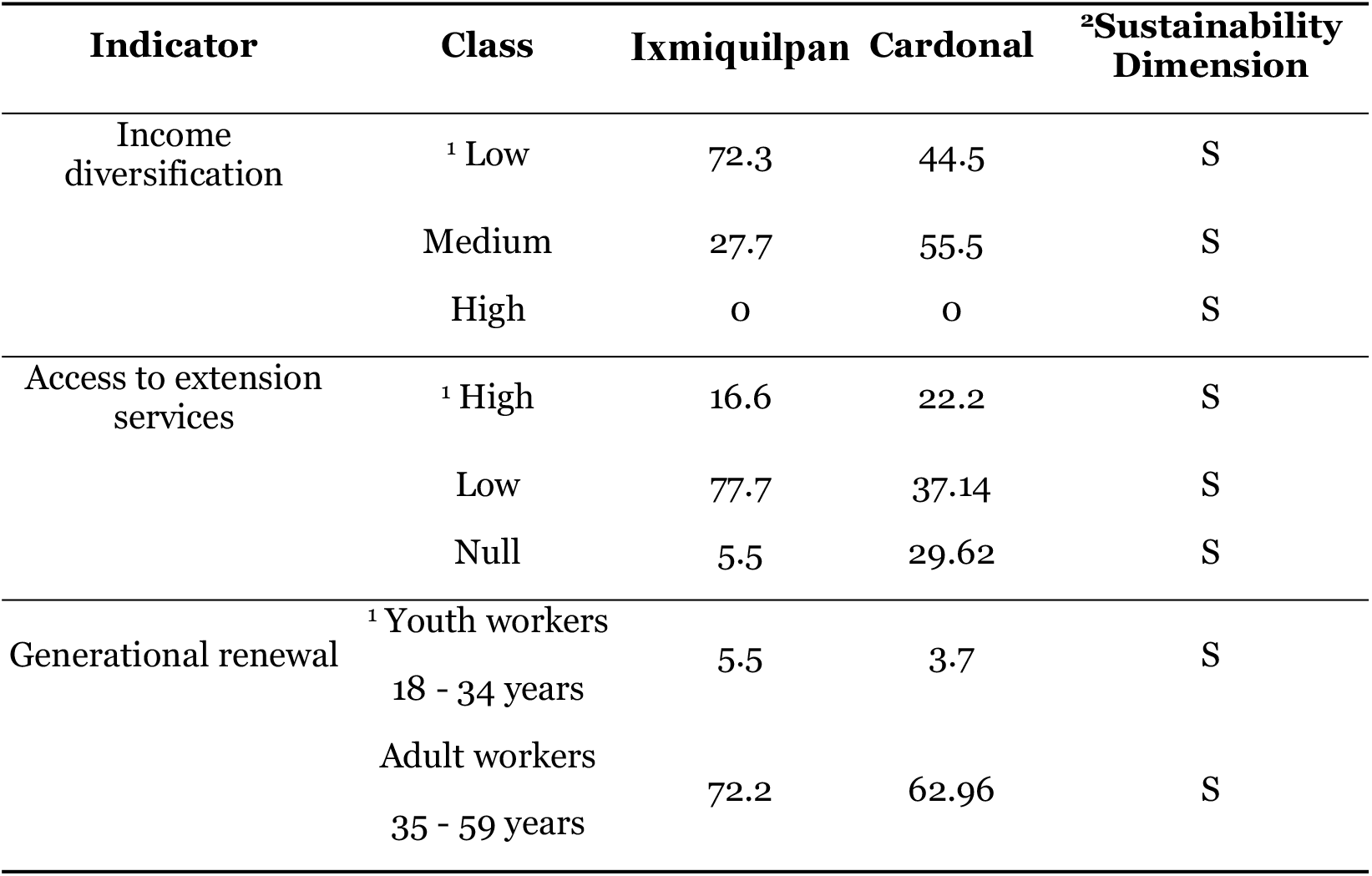
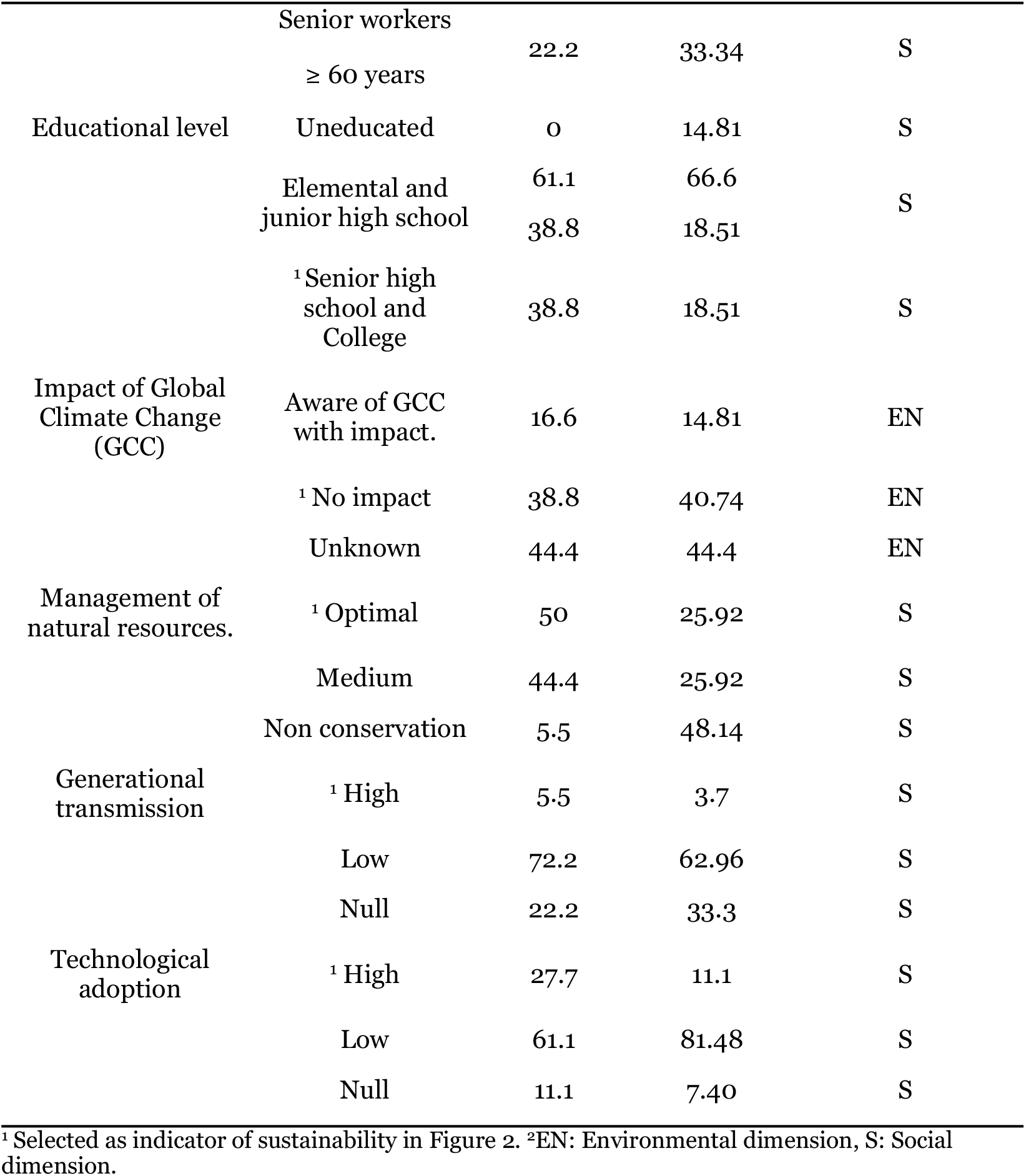
Adaptability indicators for El Cardonal and Ixmiquilpan municipalities.

#### 3.2.4. Equity

Equity focuses mainly on social justice and profit distribution. The Gini coefficient indicates an inequity of income distribution, with a value of 0.26 for Ixmiquilpan and 0.44 for El Cardonal. In Ixmiquilpan this coefficient was the same for irrigation and rainfed systems, but it was different in El Cardonal, the values were 0.24 for irrigated lands and 0.46 for rainfed. Indeed, since the profit equity is higher in El Cardonal for rainfed agriculture, the introduction of irrigation systems could have a negative impact on equity (Table 6)

**Table 6.** Equity indicators for El Cardonal and Ixmiquilpan municipalities.

| Indicator | Class | Ixmiquilpan | Cardonal | <sup>2</sup>Sustainability Dimension |
| --- | --- | --- | --- | --- |
| Infrastructure | <sup>1</sup> High | 17.1 | 18.51 | E |
|  | Medium | 68.5 | 44.4 | E |
|  | Null | 14.2 | 37.03 | E |
| Agricultural advisory services | Request | 57.14 | 62.96 | S |
|  | <sup>1</sup>Does not request | 40 | 33.3 | S |
|  | Null | 2.8 | 3.7 | S |
| Permanent workforce | <sup>1</sup>Family members per ha | 1.43 | 1.95 | S |
| Demand for temporary workforce | <sup>1</sup> Wages per ha | 2.16 | 2.29 | S |
| Distribution of machinery and vehicle | <sup>1</sup>Own (tractor) | 15.6 | 7.4 | E |
|  | <sup>*</sup>Borrowed (tractor) | 5.5 | 3.7 | E |
|  | <sup>1</sup>Rented (tractor) | 61.1 | 77.7 | E |
|  | Null (tractor) | 16.6 | 11.1 | E |
|  | Own (vehicle) | 77.7 | 66.6 | E |
|  | Borrowed (vehicle) | 0 | 0 | E |
|  | Rented (vehicle) | 0 | 0 | E |
|  | Null (vehicle) | 22.2 | 33.3 | E |
| Income distribution (Gini coefficient) | <sup>1</sup> Total agricultural activity | 0.2623 | 0.4455 | E |
|  | Irrigation system | 0.2296 | 0.2494 | E |
|  | Rainfed system | 0.2234 | 0.4681 | E |
| Women participation | <sup>1</sup> Woman | 27.7 | 18.51 | S |

<sup>1</sup> Selected as indicator of sustainability in Figure 2. <sup>2</sup>E: Economic dimension, S: Social dimension.

Women work in agriculture in both municipalities; indeed, 27 percent of farmer are women in Ixmiquilpan and 18 percent in El Cardonal (Table 6). However, women still face cultural and legal discrimination, such as lack of access to land, financing, markets, agricultural training, and education; as well as suitable working conditions, and equal treatment, therefore they are at disadvantage. Migration of youth male have given girls the opportunity to access to higher education levels, and thus they are on the way of empowering.

#### 3.2.5. Self-management and self-sufficiency

The autonomy of farmers for controlling their crops and household economy refers to self-management and self-sufficiency. The farmer’s decision to use external agricultural supplies depends on the price and availability of seeds, agrochemicals, and other products with local suppliers. Everyone purchases by himself, usually at a very high prices, what he or she can find, since they are not organized. In Ixmiquilpan, more than half of the farmers use chemical fertilizers, and in El Cardonal only 11.1 percent. However, in both municipalities 44.4 percent of farmers said they use pesticides (Table 7). Pesticides in El Cardonal are used for control of pest in agaves rather than on staple crops, while in Ixmiquilpan pest control by fumigation and use of chemical fertilizers is a common practice in irrigated systems (Table 7).

**Table 7.**
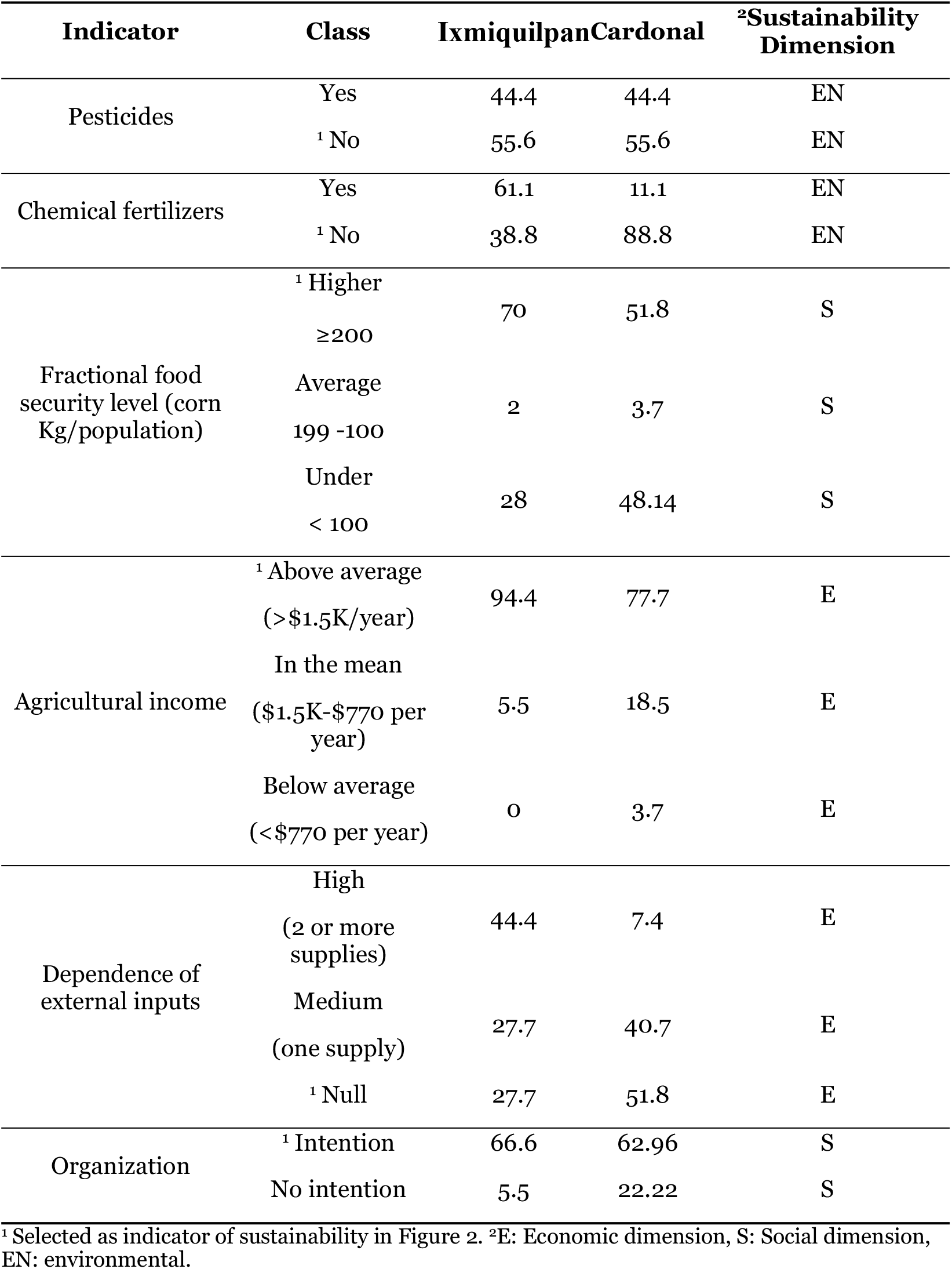
Self-management and self-sufficiency for El Cardonal and Ixmiquilpan municipalities.

Corn is the main food staple in Mexico, Mexicans consume an average of 200 Kg per person per year [27]. The corn-fractional food self-security (i.e., the extent to which the farmers can satisfy the corn needs of their families from their own crops) is sufficient for 51% of farmers in El Cardonal and 70% in Ixmiquilpan(Table 7). Thus, drought may have a higher effect on food security in El Cardonal than in Ixmiquilpan.

#### 3.2.6. Radar chart

The indicators selected from Tables 3 to 7 were used to identify the critical points of each attribute in the radar chart (Figure 2). Each indicator was plotted on a relative scale from 0 to 100; the optimal value for sustainability expected for the radar chart is 100. The adaptability is minimal for both municipalities, since all indicators are below 40 percent, excepting income diversification, because famers realize several activities besides agriculture. Stability-resilience-reliability attribute has two indicators near to optimal value, these are corn seed variety and fertilizers for El Cardonal and the last one for Ixmiquilpan. This optimal value for corn seed variety is associated to native corn seeds cultivated in the rainfed systems of El Cardonal. On the other side, the farmers of Ixmiquilpan use commercial seeds in irrigated systems and native varieties in their backyards. Solid livestock wastes are handled as fertilizer in both municipalities. There is an important difference between both municipalities on stability-resilience-reliability attribute, this difference is due to the use of commercial varieties of crops and fruit trees in Ixmiquipan and use of native seeds and native varieties in El Cardonal.

**Figure 2.**
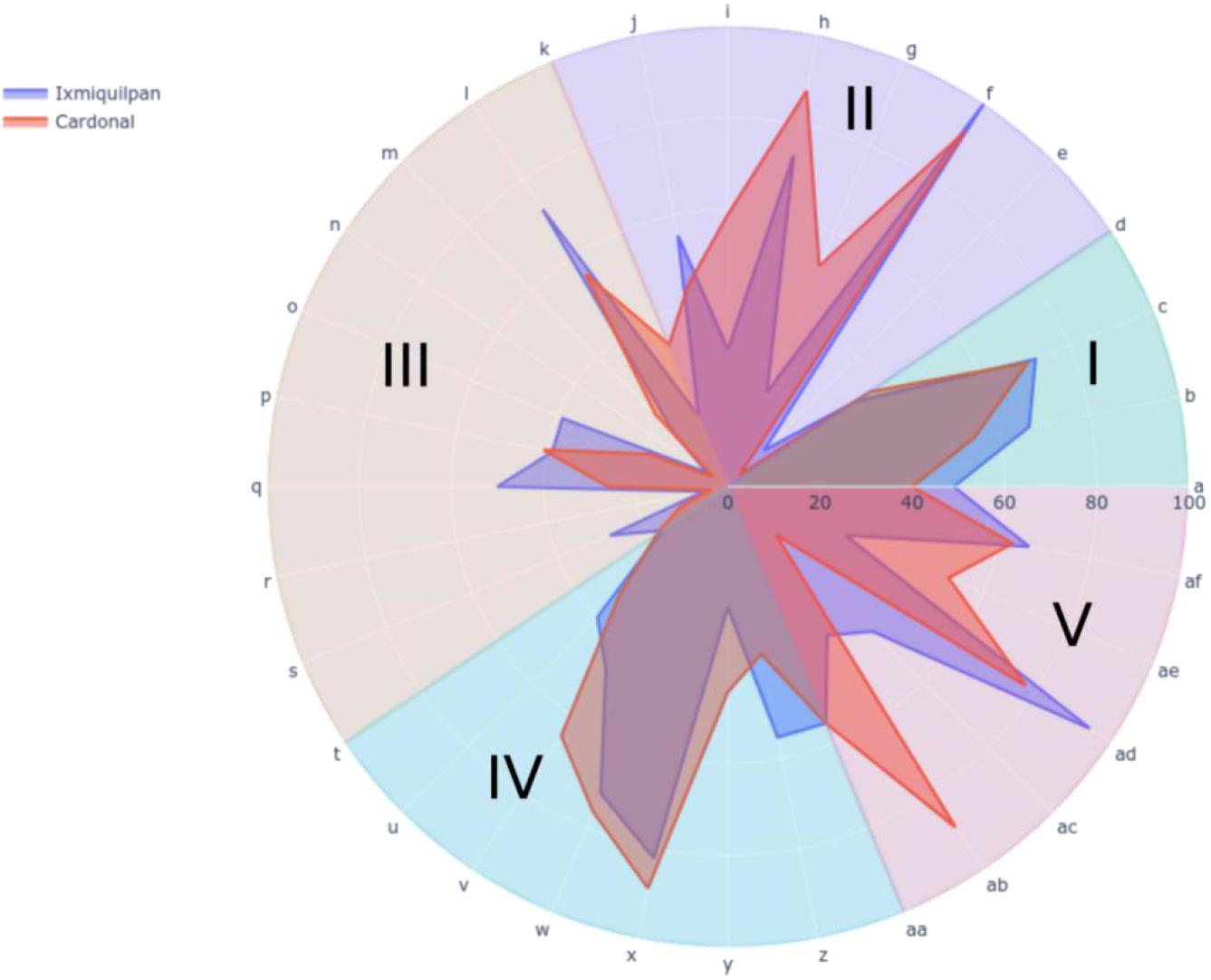
Indicators of sustainability for Ixmiquilpan and El Cardonal municipalities. Each color sector corresponds to an attribute, starting in indicator “a” and continuing in an anticlockwise direction. Attributes are: (I) productivity, (II) Stability, resilience, and reliability, (III) Adaptability, (IV) Equity, and (V) Self-management and self-sufficiency. The indicators are: (a) Agricultural total sales (irrigation), (b) Agricultural total sales (rainfed), (c) Cost-Benefit Ratio(CBR), (d) Agricultural supplies, (e) Water access, (f) Fertilizers, (g) Agrobiodiversity, (h) Corn Seed variety, (i) Seed variety of other crops, (j) Agroecological practices, (k) Pests and diseases, (l) Income diversification, (m) Access to extension services, (n) Generational renewal, (o) Educational level, (p) Impact of Global Climate Change (GCC), (q) Management of natural resources, (r) Knowledge Generational knowledge transfer, (s) New technology adoption, (t) Infrastructure, (u) Agricultural advisory services, (v) Permanent workforce, (w) Demand for temporary workforce, (x) Distribution of machinery and vehicles, (y) Income distribution (Gini coefficient), (z) Women participation, (aa) Pesticides, (ab) Chemical fertilizers, (ac) Corn-fractional food security level (corn Kg/population), (ad) Income from Agriculture, (ae) Dependence of external inputs, (af) Organization.

For the next attributes, three indicators of equity are over 50% (permanent workforce, distribution of machinery and vehicles, and temporal workforce) in both municipalities. Related to external inputs; El Cardonal has a dependence of less than 50 percent while it is 73 percent for Ixmiquilpan. In the case of self-management, the most relevant indicator for Ixmiquilpan is the average agricultural income, with a value of 92 percent, while for El Cardonal it was the minimal use of chemical fertilizers. On the other hand, the organization indicator was similar for both municipalities, as well as the cost-benefit ratio, although the total agricultural sales were slightly higher for Ixmiquilpan. (Figure 2).

Since rainfall is minimal in both municipalities (lower than 150 mm per year), water availability is critical for agriculture and thus for stability, resilience, and reliability (Fig 2). This indicator is higher in El Cardonal than in Ixmiquilpan because the water sources are rainfall and underground water (7% of water availability [29]) in the first and wastewater from Mexico City in the last.

## 4. Discussion

The evaluation of sustainability through the MESMIS methodology involves different attributes, which allow us to design strategies and identify critical points for the implementation of programs for agricultural sustainability. In both municipalities, the agricultural sales are low, but the total incomes are high; therefore, the cost-benefit is in a medium range. This discrepancy is due to migrant remittances.

The critical points identified in both municipalities belong to attributes adaptability, equity, and self-management and self-sufficiency. All these attributes indicate an economic and food security fragility, associated to the inequity of income distribution and low corn-fractional food security (Figure 2.). Fragility is understood as the vulnerability of small farm holders to changes or uncertainty in the economic, social, and environmental conditions [30]. The external factor migration has an important impact on both economic and food security fragility (Figure 3), indeed, in 2015, injection of remittances represented from 9.0 to 14.6% of the gross domestic product of Ixmiquilpan and 14.6 to 36.2% in El Cardonal [31].

**Figure 3.**
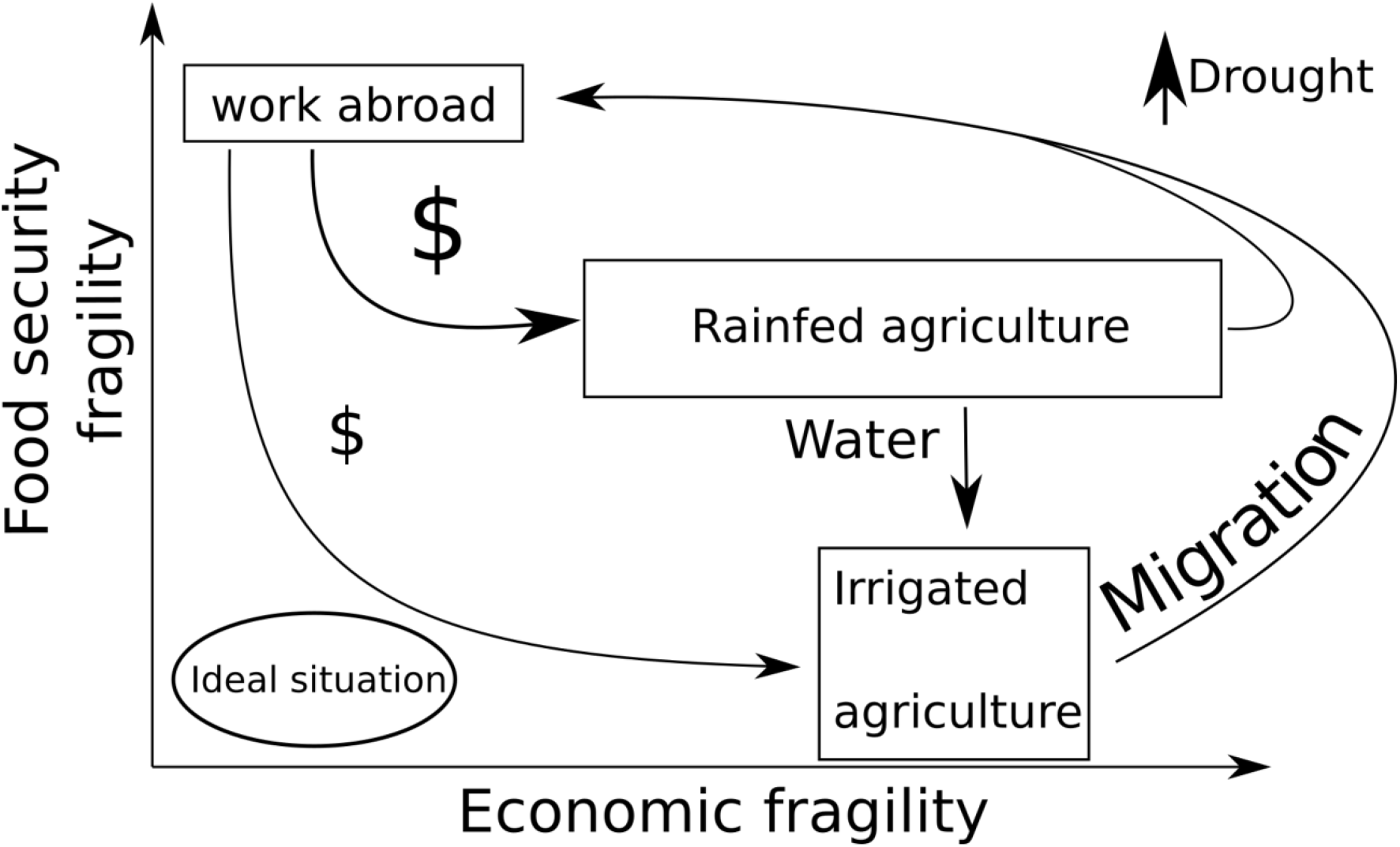
Model of interactions between economic and food security fragilities.

The small farm holders in Ixmiquilpan depend on external agricultural supplies and their agricultural infrastructure is scarce, but the remittances are not use for improving this infrastructure neither, for agriculture innovation or for acquiring supplies; remittances are mainly use for purchased food, for satisfying other necessities as well as building houses for the family. Thus remittances lead to an unsustainable economic cycle, increasing the fragility of the agricultural system (Figure 3). On the other hand, in El Cardonal, money injection does not compensate the low agriculture yields and the producers must diversify their activities to be able to satisfy their food necessities, and as a result they neglect their farmlands. Thus, remittances increase the vulnerability of agriculture in a long time. Furthermore, the increase of food security fragility is affected by insufficiency of water, decrease of native corn consumption and increment of industrialized food products. Previous studies developed on mesquite tree (*Prosopis laevigata Humb. et Bonpl. ex Willd*) in a community of Ixmiquilpan [32], also pointed to the influence of migration on the stability and reliability attributes, on the lack of generational knowledge transmission that affects the adoption of traditional techniques by new generations.

In fact remittances may have positive and negative effects in Mexican rural communities. Edward Taylor et. al. studied the impact of migrant remittances on the distribution of rural income and rural poverty in Mexico [33]. They found that remittances from migrants abroad slightly increase rural income inequalities, while remittance of internal migrants are income equalizer, but in high migration regions at long time both type of remittances have an equalizing effect in income. Ixmiquipan is a high migration municipality and migration is even higher in El Cardonal, therefore remittance may have an equalizer effect on income. Hikmet Ersek, President and Chief Executive Officer of Western Union said, remittances generate crucial positive, economic and social effects in developing countries. However, we found that remittances in agricultural systems in long term reduces the likelihood of the family members to continue agricultural activities, they also have a negative effect in cultural identity, and in the traditional diet of indigenous people. Indeed, remittances decreases the possibility to transfer the traditions and cultural practices to young generations, affects the adoption of new technologies and the appropriation of new solutions to improve agriculture yields, water resource management and sustainability. Thus, abroad migration remittances have at least two faces.

On the other hand, the use of untreated wastewater from Mexico City in the agriculture irrigation systems impacts agriculture both in positive and negative ways. On one side, it has a negative effect on the inhabitants’ health and increases the vulnerability to climate change [34]. Indeed, the increment of agricultural yields due to wastewater irrigation is linked to a high investment on supplies, unsafe food products, diseases and malnutrition, and finally it affects the economic income [35]. The social problems related to health are critical for the development of dry lands. On the other side, watewater has in a short term, a positive effect in agricultural yields. However, in the environmental context there are other options for using and regenerating dry lands, for example, Luján Soto et al. recommend a combination of techniques [36], such as the reforestation of local crops, like agave or cactus and technical services for control of plagues and phytopathogens. In this context the adoption of climate smart agriculture has impacts in carbon sequestration and biodiversity conservation; in fact, it could generate additional incomes for the small holders by ecosystem services [37].

In summary: 1) Adaptability is a critical factor in both municipalities and it does not depend on economic factors, rather the main problem is youth migration. 2) Access to water, and economic resources as well as management of environmental resources are an imperious need to increase the yield of agriculture crops and equity. 3) Strengthening of resilience requires organization of small producers and the combination of indigenous and modern technologies for territorial development. 4) The social adaptation of both communities is in a critical level, because of the generational brake between farmers and their offspring. 5) Productivity depends mostly on agricultural supplies affecting farmers CBR that is similar for rainfed and irrigation agriculture. 6) Remittances have two faces, a positive effect on income and poverty reduction, and a negative one on agriculture, traditional diet and cultural identity(Figure 2, Figure 3).

## 5. Conclusions

The vulnerability of the agricultural systems to crop pests and diseases needs to be addressed, as it reduces productivity and the associated profits, forcing peasants to seek other economic activities. Furthermore, technical services are also needed to apply other agroecological practices for solving problems such as optimal management of water resources and pest and diseases control. The lack of technical services, together with the economic problems, contribute to breaking the generation renewal, and force the youth to migrate to find a better life.

Some recommendations for both municipalities, to reduce the food security fragility, are implementation of integral systems for water management, including rainfall capture, underground water, and treated wastewater, as well as the implementation of efficient irrigation systems for decreasing water and energy consumptions. These innovations should integrate traditional knowledge and modern science and technology including producers, youths, men, and women. The organization of small farm holders is crucial for the development of long-term sustainability projects.

Other long-term sustainable recommendations are: 1) promotion of polycultures in the region using native staple crops to increase food security, and lead to a robust economic income; 2) promotion of rural sustainable enterprises, to increase te values of agricultural products, organizations of producers and their familes must own these companies, 3) appropriation by producers of the supply chain and distribution channels.

These recommendations are aimed at increasing the economic sustainability of agriculture and its resilience to climate change. More robust and profitable crops would be more attractive to young farmers, stimulating the cross-generational continuity. In that context, migration would turn from a necessity to an option for youths. Also, more productive agricultural systems would encourage farmers and their relatives to invest in agriculture. Thus, by strengthening the agriculture system, the fragility of the socio-economic system of the region would be improved overall, with less migration, and more regional benefits from. Finally, a strengthened agriculture could bring robustness for the region in the case of macroeconomic disturbances, as it would ensure food security, in terms of self-nourishment.

## Author Contributions

Conceptualization, Y.C. and D.L.; methodology, Y.C.; validation, Y.C., D.L. and M.D.; formal analysis, Y.C.; investigation, Y.C.; resources, D.L.; data curation, M.D., Y.C.; writing—original draft preparation, Y.C.; writing—review and editing, M.D., D.L., Y.C.; visualization, Y.C.; funding acquisition, D.L. All authors have read and agreed to the published version of the manuscript.

## Data Availability Statement

The data presented in this study are available on request from the corresponding author.

## Acknowledgments

The authors would like to thank E Aguirre von Wobeser and Pacblo Wong for the valuable recommendations. Inhabitants of Ixmiquilapan and El Cardonal municipalities for their willingness. To the team of the Regional Unit Hidalgo of Centro de Investigacion en Alimentacion y Desarrollo A. C. for the recollection of data.

## References

1. FAO. Dimensions of need - Sustainable agriculture and rural development http://www.fao.org/3/u8480e/U8480E0l.htm (accessed May 31, 2021).

2. Tommasino, H. Sustentabilidad rural: desacuerdos y controversias. In ¿Sustentabilidad?: desacuerdos sobre el desarrollo sustentable; Porrua: Montevideo, Argentina, 2001; pp 139–163.

3. Foladori, G.; Tommasino, H. El concepto de desarrollo sustentable treinta años después. Desenvolvimento e Meio Ambiente 2000, 1 (0). https://doi.org/10.5380/dma.v1i0.3056.

4. Bejarano Ávila, J. A.; Aldana Navarrete, H.; Meek Muñoz, E.; Agricultura (IICA), I. I. de C. para la. Desarrollo sostenible: un enfoque económico con una extensión al sector agropecuario; 1998.

5. Andreoli, M.; Tellarini, V. Farm Sustainability Evaluation: Methodology and Practice. Agriculture, Ecosystems & Environment 2000, 77 (1), 43–52. https://doi.org/10.1016/S0167-8809(99)00091-2.

6. Valdez-Vazquez, I.; del Rosario Sánchez Gastelum, C.; Escalante, A. E. Proposal for a Sustainability Evaluation Framework for Bioenergy Production Systems Using the MESMIS Methodology. Renewable and Sustainable Energy Reviews 2017, 68, 360–369. https://doi.org/10.1016/j.rser.2016.09.136.

7. Astier, M.; García-Barrios, L.; Galván-Miyoshi, Y.; González-Esquivel, C.; Masera, O. Assessing the Sustainability of Small Farmer Natural Resource Management Systems. A Critical Analysis of the MESMIS Program (1995-2010). Ecology and Society 2012, 17 (3). https://doi.org/10.5751/ES-04910-170325.

8. van der Werf, H. M. G.; Petit, J. Evaluation of the Environmental Impact of Agriculture at the Farm Level: A Comparison and Analysis of 12 Indicator-Based Methods. Agriculture, Ecosystems & Environment 2002, 93 (1), 131–145. https://doi.org/10.1016/S0167-8809(01)00354-1.

9. Romero-Arenas, O.; Morales, P. S. Evaluación de la sustentabilidad del sistema milpa en el estado de Tlaxcala, México. Revista de El Colegio de San Luis 2018, No. 15, 10.

10. Talukder, B.; Blay-Palmer, A. Comparison of Methods to Assess Agricultural Sustainability. In Sustainable Agriculture Reviews; Lichtfouse, E., Ed.; Sustainable Agriculture Reviews; Springer International Publishing: Cham, 2017; pp 149–168. https://doi.org/10.1007/978-3-319-58679-3_5.

11. Ixmiquilpan Monthly Climate Averages https://www.worldweatheronline.com/ixmiquilpan-weather/hidalgo/mx.aspx (accessed May 25, 2021).

12. Cardonal Monthly Climate Averages https://www.worldweatheronline.com/cardonal-weather/hidalgo/mx.aspx (accessed May 25, 2021).

13. Mexico’s Indigenous Population http://www.culturalsurvival.org/publications/cultural-survival-quarterly/mexicos-indigenous-population (accessed Jun 2, 2021).

14. Institute for Federalism and Municipal Development (INAFED). Ixmiquilpan. Encyclopedia of the Municipalities and Delegations of Mexico. Ministry of the Interior of Mexico; Institute for Federalism and Municipal Development (INAFED), 2010. http://www.inafed.gob.mx/work/enciclopedia/EMM13hidalgo/municipios/13030a.html

15. Encuesta Nacional Agropecuaria 2019 https://www.inegi.org.mx/programas/ena/2019/ (accessed May 14, 2021).

16. Secretary of Social Development (SEDESOL). Hidalgo, Cardonal: Annual Report on the Situation of Poverty and Social Backwardness 2017. Undersecretary of Planning, Evaluation and Regional Development. SEDESOL. Secretary of Social Development (SEDESOL) 2017.

17. Secretary of Social Development (SEDESOL). Ixmiquilpan: Annual Report on the Situation of Poverty and Social Backwardness 2017. Undersecretary of Planning, Evaluation and Regional Development. SEDESOL. Secretary of Social Development (SEDESOL) 2017.

18. KoBoToolbox | Data Collection Tools for Challenging Environments https://kobotoolbox.org/ (accessed Mar 4, 2021).

19. Jr, K. P. N.; Grimm, L. G. Statistical Applications for the Behavioral and Social Sciences; John Wiley & Sons, 2018.

20. Elementary Statistics, 13th Edition /content/one-dot-com/one-dot-com/us/en/higher-education/program.html (accessed Mar 4, 2021).

21. UNESCO y UNAM, CIGA Centro de Investigaciones en Geografía Ambiental de la Universidad Nacional Autónoma de México. Sostenibilidad en sistemas de manejo de recursos naturales en países andinos, 1st ed.; IGA Centro de Investigaciones en Geografía Ambiental de la Universidad Nacional Autónoma de México: México, 2018; Vol. 1.

22. López-Ridaura, S.; Masera, O.; Astier, M. Evaluating the Sustainability of Complex Socio-Environmental Systems. the MESMIS Framework. Ecological Indicators 2002, 2 (1), 135–148. https://doi.org/10.1016/S1470-160X(02)00043-2.

23. Speelman, E. N.; López-Ridaura, S.; Colomer, N. A.; Astier, M.; Masera, O. R. Ten Years of Sustainability Evaluation Using the MESMIS Framework: Lessons Learned from Its Application in 28 Latin American Case Studies. International Journal of Sustainable Development & World Ecology 2007, 14 (4), 345–361. https://doi.org/10.1080/13504500709469735.

24. Altieri, M. A.; Nicholls, C. I.; Henao, A.; Lana, M. A. Agroecology and the Design of Climate Change-Resilient Farming Systems. Agron. Sustain. Dev. 2015, 35 (3), 869–890. https://doi.org/10.1007/s13593-015-0285-2.

25. Farris, F. A. The Gini Index and Measures of Inequality. The American Mathematical Monthly 2010, 117 (10), 851–864. https://doi.org/10.4169/000298910X523344.

26. Witlox, F. Gini Coefficient. In International Encyclopedia of Geography; American Cancer Society, 2017; pp 1–4. https://doi.org/10.1002/9781118786352.wbieg0855.

27. SAGARPA. MAÍZ GRANO BLANCO Y AMARILLO Mexicano; Planeación Agrícola Nacional 2017-2030; Secretaria de Agricultura, Ganaderia, Desarrollo Rural, Pesca y Alimentación: México, 2018; p 28.

28. INEGI. Panorama sociodemográfico de Hidalgo. Censo de Población y Vivienda 2020, 1st ed.; Panorama sociodemográfico de; INEGI: México, 2021; Vol. Hidalgo.

29. CONAGUA. ACTUALIZACIÓN DE LA DISPONIBILIDAD MEDIA ANUAL DE AGUA EN EL ACUÍFERO VALLE DEL MEZQUITAL(1310) ESTADO DE HIDALGO; DISPONIBILIDAD MEDIA ANUAL DE AGUA EN EL ACUÍFERO VALLE DEL MEZQUITAL; Actualización 1310; CONAGUA: Hidalgo, Mexico, 2020; p 42.

30. Taleb, N. N. Antifragile: Things That Gain from Disorder; 2016.

31. Cortés Rivera, D.; Granados Alcantar, J. A.; Quezada Ramírez, M. F. International Migration in Hidalgo: News Dynamics and Actors. Economía, sociedad y territorio 2020, 20 (63), 429–456. https://doi.org/10.22136/est20201557.

32. Pérez-Serrano, D.; Cabirol, N.; Martínez-Cervantes, C.; Rojas-Oropeza, M. Mesquite Management in the Mezquital Valley: A Sustainability Assessment Based on the View Point of the Hñähñú Indigenous Community. Environmental and Sustainability Indicators 2021, 10, 100113. https://doi.org/10.1016/i.indic.2021.100113.

33. Taylor, J. E.; Mora, J.; Adams, R.; Lopez-Feldman, A. Remittances, Inequality and Poverty Evidence from Rural Mexico. In Migration and Development Within and Across Borders: Research and Policy Perspectives on Internal and International Migration; International Organization for Migration and The Social Science Research Council, 2008; pp 101–130.

34. Durán-Álvarez, J. C.; Jiménez, B.; Rodríguez-Varela, M.; Prado, B. The Mezquital Valley from the Perspective of the New Dryland Development Paradigm (DDP): Present and Future Challenges to Achieve Sustainable Development. Current Opinion in Environmental Sustainability 2021, 48, 139–150. https://doi.org/10.1016/j.cosust.2021.01.005.

35. FAO - News Article: Exploring the use of wastewater in agriculture http://www.fao.org/news/story/en/item/463433/icode/ (accessed May 25, 2021).

36. Luján Soto, R.; Martínez-Mena, M.; Cuéllar Padilla, M.; de Vente, J. Restoring Soil Quality of Woody Agroecosystems in Mediterranean Drylands through Regenerative Agriculture. Agriculture, Ecosystems & Environment 2021, 306, 107191. https://doi.org/10.1016/j.agee.2020.107191.

37. Sain, G.; Loboguerrero, A. M.; Corner-Dolloff, C.; Lizarazo, M.; Nowak, A.; Martínez-Barón, D.; Andrieu, N. Costs and Benefits of Climate-Smart Agriculture: The Case of the Dry Corridor in Guatemala. Agricultural Systems 2017, 151, 163–173. https://doi.org/10.1016/j.agsy.2016.05.004.

